# Comprehensive Mapping of Human dsRNAome Reveals Conservation, Neuronal Enrichment, and Intermolecular Interactions

**DOI:** 10.1101/2025.01.24.634786

**Authors:** Ryan J. Andrews, Brenda L. Bass

## Abstract

The human transcriptome contains millions of A-to-I editing sites arising from an unclear number of poorly characterized dsRNAs. Editing sites are often used to infer presence of dsRNA, but this method is limited by transcription levels, read depth, ADAR expression and cannot identify unedited dsRNA. To address these limitations, we developed dsRNAscan. Applying dsRNAscan to the human genome predicted 5 million dsRNAs. Genomic distribution of dsRNAs encompassing repetitive elements was widespread, but non-repetitive dsRNAs were sparse and enriched at chromosomal tips. dsRNAscan predicted hundreds of long, highly paired dsRNAs suspected to be immunogenic, but only one was in a 3′UTR, and thus likely to challenge cytoplasmic immune sensors. We observed several thousand editing enriched regions suspected to arise from intermolecular structures, and dozens of neuronally enriched dsRNAs conserved across vertebrates. This study offers the first comprehensive set of dsRNA annotations for the human genome, available as a resource at https://dsrna.chpc.utah.edu/.

## INTRODUCTION

RNA sequencing-based methods are used to infer A-to-I editing sites,^1^ which are catalyzed by adenosine deaminases acting on RNA (ADARs)—enzymes that exclusively bind and edit dsRNA helices.^2^ A-to-I editing analyses reveal that the human genome encodes a surprising amount of dsRNA, serving as the substrate for over 15 million editing sites.^3,4^ The presence of ADAR editing sites is the gold standard for positively identifying dsRNA that exists *in vivo* and has been used across multiple organisms.^5–7^ However, RNA-seq based methods have limitations: they are dependent on transcription, read depth, and/or sufficient ADAR expression levels.^8^ Further, using an approach *based on editing* cannot identify unedited dsRNA. Consequently, the dsRNA-forming potential of RNAs that are lowly transcribed or in tissues with low ADAR expression, remains unknown. Further, only a limited number of editing sites have undergone secondary structure predictions,^6,9,10^ so it is unclear how structurally diverse RNA editing substrates are; for example, how often do they contain bulges, mismatches, and internal loops, and how rare are perfectly paired dsRNA? To address these outstanding questions, we devised dsRNAscan, a sequence-based, editing-neutral, computational approach to predict dsRNA formation.

We utilized dsRNAscan to analyze the human genome, predicting over five million putative dsRNAs containing at least 25 base pairs (bps). Each dsRNA was evaluated for compositional and thermodynamic properties using custom algorithms and tools available in the ViennaRNA package.^11^ Intersecting predicted dsRNAs with various genome annotations allowed for in-depth categorization of all dsRNA predictions. Our results confirmed that most dsRNA is edited and formed between inverted repetitive elements, especially Alu sequences. Conversely, prediction of diverse *unedited* dsRNAs, provided insight into structural properties that disfavor ADAR editing. These differences allowed for development of a machine learning model to classify dsRNAs based on their likelihood of being ADAR substrates, providing a robust tool for future dsRNA prediction and analysis. The genomic distribution of dsRNAs revealed widespread repetitive dsRNAs, while non-repetitive dsRNAs were sparse but enriched at chromosomal tips. In the search for immunogenic dsRNA, dsRNAscan identified approximately 2,000 dsRNAs that are likely to be immunogenic and sensed by MDA5, the primary sensor in autoimmune responses to endogenous dsRNA.^12,13^ However, notably, only *one* of these was located in a 3′UTR (in the gene ZNF426), and thus predicted to be accessible to cytoplasmic MDA5. Additionally, we investigated the potential for intermolecular dsRNA formation, which has been implicated as an alternative substrate for MDA5 in several studies,^14,15^ and identified several thousand editing-enriched regions likely originating from such intermolecular structures. Finally, we identified several hundred conserved dsRNAs that are enriched in genes highly expressed in the cerebral cortex and involved in neuronal development, emphasizing dsRNA’s importance in brain function.

Results have been converted into standardized file formats including base pair (.bp) files that can be used to easily visualize span and magnitude of dsRNA as large arcs on genome browsers,^16^ and GFF3/BED files that can be used to analyze dsRNAs alongside a multitude of existing genomic datasets. These analyses represent the first comprehensive set of dsRNA annotations for the human genome and are available for download and shared directly via an Integrated Genome Browser web app (https://dsrna.chpc.utah.edu/).

## RESULTS

### dsRNAscan predicted over five million dsRNAs

dsRNAscan was constructed in Python and utilizes C++ from the EMBOSS package^17^ and RNAduplex from the ViennaRNA suite^11,18,19^ to predict dsRNA across the genome (**Figure 1A**; see Methods). First, dsRNAscan fragments each strand of the genome into overlapping analysis windows spanning 10,000 nucleotides (nts); the selection of this window size was informed by studies showing the majority of long-range RNA *intra*molecular interactions in the human transcriptome span less than 10,000 nts.^20,21^ Successive windows were created after stepping forward 150 nts until each strand of the genome was covered by over 20 million overlapping windows. Within each window, the einverted algorithm first identified sequences capable of forming dsRNA structures based on the number of bps, mismatches, and gaps. The coordinates of each arm of the dsRNA is recorded, with the loop implicitly defined by the coordinates of the region between each arm (**Figure 1A**). Importantly, we modified einverted to recognize GU mismatches as positively scoring matches, since these thermodynamically favorable bps are often found in RNA structures. With GU pairs scoring positively, we used a score threshold of 75 to maintain stringent detection and avoid excessive false positives (**see Methods**), whereby the score was calculated based on number of base pairs (matches), internal loops (mismatches), and bulges (gaps), scoring +3, -4, and -12 respectively (**Figure 1B)**. This effectively limited our search to dsRNAs with at least 25 bps, the minimum number of base pairs required to reach a score of 75. Mathematically then, bulges, mismatches, and internal loops were allowed only for dsRNAs longer than 25 base pairs. Following this, RNAduplex was used to estimate the thermodynamic stability of all einverted results, construct dot bracket notation structures,^22^ and base pair percentages. As a final filter, any prediction with less than 70% base pairs was discarded. This approach, informed by observations that RNA transcription is pervasive,^23,24^ was designed to map the full potential of dsRNA formation encoded in the transcribed human genome. Implementing this across the human genome resulted in the prediction of 5.1 million dsRNAs (**Table S1;** https://dsrna.chpc.utah.edu/shiny/dsrna/).

**Figure 1:**
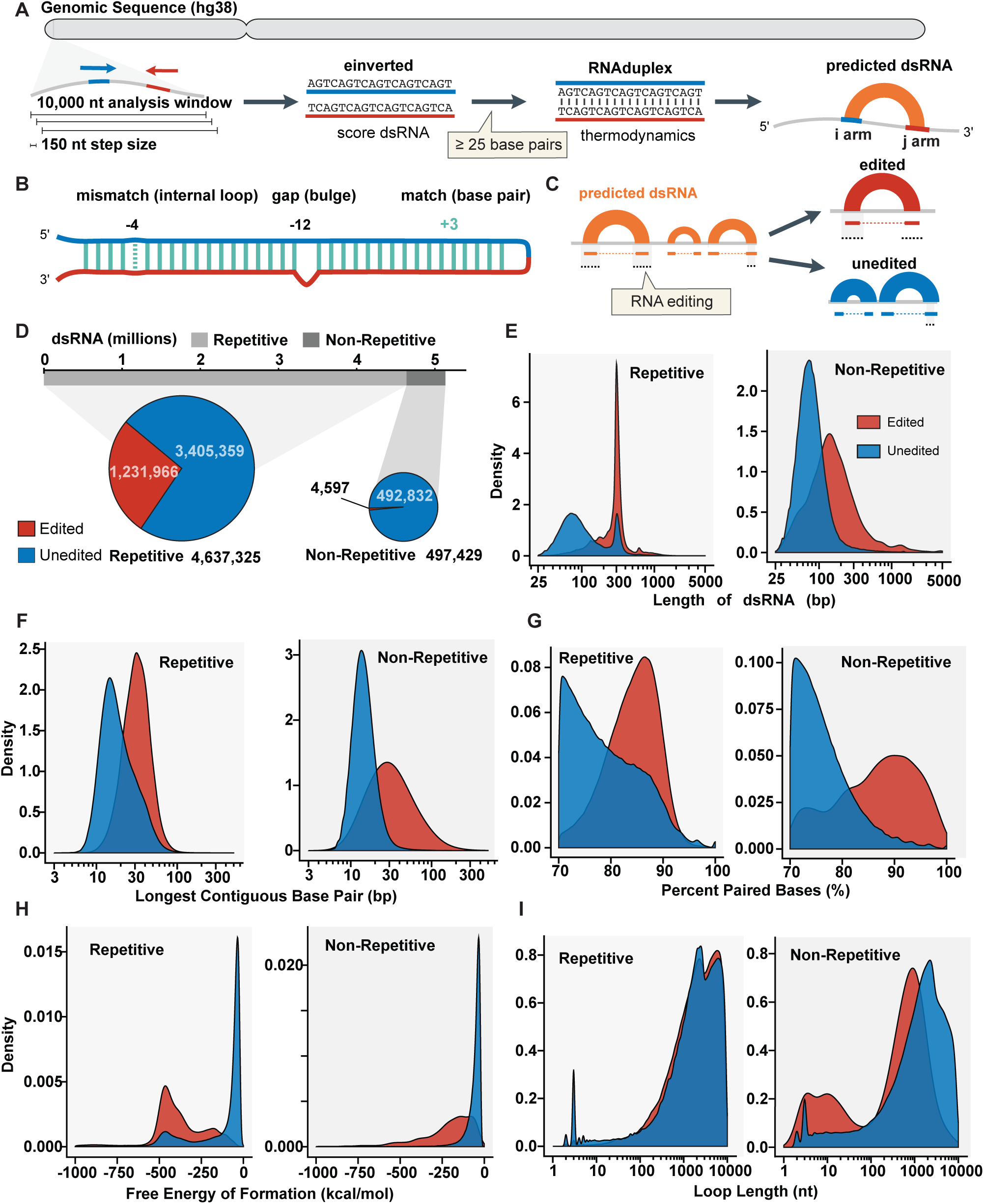
Overview of dsRNAscan and its results for the human genome. **A.** Schematic of dsRNAscan pipeline. Genomic sequences are divided into analysis windows and scored with einverted to detect inverted repeats capable of forming dsRNA. RNAduplex is then used to evaluate thermodynamic stability, with resulting predicted dsRNAs depicted as orange arches connecting arms of dsRNA. **B.** Scoring criteria for predicted dsRNAs: matches (+3), mismatches/internal loops (-4), and gaps/bulges (-12). Only dsRNAs with a score ≥ 75 are selected, ensuring ≥ 25 base pairs are present, with allowances for mismatches, which correspond to internal loops of various sizes, and gaps, which structurally correspond to bulges on opposite strand. **C.** Schematic of RNA editing, showing how predicted dsRNAs are classified as edited if editing sites occur on both arms. **D.** Pie chart showing total number of predicted dsRNAs, divided into repetitive and non-repetitive categories, with subcategories of edited and unedited dsRNAs. **E.** Density plots of dsRNA length distribution for non-repetitive and repetitive dsRNAs. The y-axis represents the kernel density estimate, which gives a smoothed approximation of frequency distribution, ensuring that area under the curve equals 1. The x-axis shows the range of dsRNA lengths. These plots allow for direct comparison of length distributions between different groups, with peaks indicating most frequent dsRNA lengths in each category. **F.** Density plots of longest contiguous base pair stretches for non-repetitive and repetitive dsRNAs, highlighting differences between edited (red) and unedited (blue) dsRNAs. **G.** Density plots of percentage of paired bases within non-repetitive and repetitive dsRNAs, comparing edited (red) and unedited (blue) groups. **H.** Density plots of free energy of formation (ΔG) for non-repetitive and repetitive dsRNAs, showing stability differences between edited (red) and unedited (blue) dsRNAs. **I.** Density plots of loop lengths in non-repetitive and repetitive dsRNAs.

### Most predicted dsRNAs are unedited

Prior genome-wide predictions of dsRNA mostly rely on the presence of ADAR editing sites, and we were interested in how our editing agnostic pipeline would compare. Accordingly, we overlaid our prediction coordinates with RNA editing coordinates from the REDIportal database which contains ∼16 million unique editing sites predicted from over 9642 RNA-seq samples^3,4^; these editing sites comprise an aggregation of editing sites from diverse tissues, and importantly, do not make any discrimination between editing from ADAR1p150, ADAR1p110 or ADAR2.^25^ Only dsRNAs containing editing sites on *both* arms were categorized as edited (**Figure 1C**). Also, since most editing in animal genomes maps to repetitive sequences,^7,26^ which comprise roughly 50% of the human genome,^27^ we further categorized dsRNAs as either “repetitive” or “non-repetitive” (**see Methods**). All dsRNA arms were intersected with repetitive elements including simple repeats, long terminal repeats, LINE retroelements, Alu retroelements and other SINE elements; dsRNA intersecting any one of these elements was categorized as repetitive, while those without any intersection were categorized as non-repetitive. While roughly 50% of the genome is composed of non-repetitive sequences, over 90% of dsRNAs and >99% of edited dsRNAs were repetitive (**Figure 1D**). The other 50% of the genome, composed of non-repetitive sequences, was only predicted to form around half a million dsRNAs with just a small number of these, 4,597, being edited (**Table S2**). These categorical results were consistent with current knowledge of RNA editing, which largely occurs in repetitive elements. For a more complete picture of the dsRNAome we next analyzed the structural characteristics of each dsRNA category.

### dsRNA structural characteristics

Short dsRNA helices are the foundation of RNA secondary structure, and these interact to form three-dimensional structures that are required for the function of many RNAs. The helices in RNA secondary structures are on average only ∼8 bps long before they are interrupted by mismatches, bulges, or loops (based on our analysis of 102,318 reference structures from bpRNA^28^; **Table S3**); in addition, there are some examples of isolated short stem-loop structures such as those involved in 3′ processing of histone mRNAs.^29^ By contrast, we were particularly interested in describing the rod-shaped dsRNA structures that typically lack tertiary interactions and are bound by dsRNA binding proteins. While such structures can be hundreds of bps long, typically at least two turns of the helix defines the minimum length for biologically relevant protein binding. For example, PKR is not robustly activated unless a dsRNA is at least ∼30 bps long,^30^ and some of the shortest but functional ADAR substrates are smaller than 30 bps long (e.g., the edited GRIA2 Q/R duplex is 28 bp long with 2 mismatches).^31^ Thus, we set minimum dsRNA length to 25 bps, with the upper limit implicitly defined by window size of 10,000 nt, which limited the maximum dsRNA length to 5000 bp, well above the length of any known intramolecular dsRNA.^32^ We found that most predicted dsRNAs were well within these two extremes: 99% of dsRNA structures were between 30 and 1000 bps long (**Figure 1E**).

In addition to length, dsRNAscan captured four other metrics to more completely describe each dsRNA: contiguous bp count, free energy of formation (ΔG), base pair percentage and loop length (**see Methods**). When considering these, we noted distinct differences between edited/unedited and repetitive/non-repetitive dsRNAs (**Table 1**; **Figure 1E**). For repetitive dsRNAs, those that encompassed editing sites were longer than those that were unedited (median length of 297 vs 87 bp respectively). Non-repetitive dsRNAs were shorter overall with unedited predictions even shorter (median length of 144 bp for edited and median length of 75 bp for unedited). Next, for each dsRNA we counted the number of contiguous, uninterrupted bps to allow comparisons by this parameter (**Figure 1F**). We found that edited dsRNAs had longer stretches of uninterrupted base pairs (median 31 vs. 17 bps), with the trend similar for repetitive and non-repetitive dsRNAs. Edited dsRNAs had a higher percentage of bps than unedited dsRNAs (**Figure 1G**; median 84.67 vs 77.42% for repetitive), and correspondingly had lower free energies (**Figure 1H**). Overall loop lengths, defined as the length of RNA intervening the two arms, were more consistent between groups at around 2000 bps on average, but there was a peak in loop lengths less than 100 nt for non-repetitive edited dsRNAs (**Figure 1I**) likely representing smaller dsRNA structures like primary miRNA stems.

**Table 1.** Summary of structural characteristics for all dsRNAs. This table contains structural metrics for edited and unedited dsRNAs across repetitive and non-repetitive regions of the human genome. Median values for key features, including dsRNA length, contiguous base pairs (bps), free energy of formation (ΔG), base-pair percentage (BP%), and loop length, are reported for both repetitive and non-repetitive dsRNAs.

|  | Total |  | Repetitive |  | Non-Repetitive |  |
| --- | --- | --- | --- | --- | --- | --- |
| dsRNA feature | Edited<br>(median) | Unedited<br>(median) | Edited<br>(median) | Unedited<br>(median) | Edited<br>(median) | Unedited<br>(median) |
| Length (bps) | 297 | 87 | 297 | 90 | 144 | 75 |
| Continuous bps | 31 | 17 | 31 | 17 | 30 | 14 |
| $\Delta G$ (kcal/mol) | -410.4 | -52.9 | -410.8 | -54.7 | -184.8 | -44.6 |
| BP (%) | 84.67 | 77.42 | 84.66 | 77.88 | 87.54 | 75.2 |
| Loop Length (nt) | 2025 | 1968 | 2035 | 2049 | 509 | 1407 |
| Number of dsRNAs | 1,236,563 | 3,898,191 | 1,231,966 | 3,405,359 | 4,597 | 492,832 |

### Machine learning predicts editing based on structural characteristics and transcription

The clear difference in properties between edited/unedited dsRNA predictions suggested we could utilize machine learning to isolate dsRNA properties that most correlated with editing. To this end, we employed the TensorFlow Decision Forests (TFDF) package^33,34^ to classify our dsRNA predictions based on their likelihood of serving as human ADAR substrates. In this application, the decision tree method predicted whether a dsRNA was edited based on a series of branching yes/no questions about its properties, allowing for insight into which properties contribute to each prediction. We developed our training set based on six core dsRNA properties: length, base pair count, loop length, percentage of bps, thermodynamic stability, and nucleotide content (including single nucleotide frequency and trinucleotide frequency which allows for consideration of ADAR nearest neighbor preferences).^35^ For this, we defined ‘true’ dsRNAs as those containing at least one editing site on both arms. To maintain the model’s generalizability for predicting structures, we tested a range of maximum depths, where higher depths allow the tree to create more branches, capturing complex patterns in the data but risking overfitting by modeling noise rather than underlying relationships. Higher tree depths (>8) increased accuracy, but were likely overfitting. When we incorporated the transcriptional status of predicted dsRNA into our model, we observed a boost in accuracy (**Table S4**). Subsequently, all dsRNAs were annotated with transcriptional data along with prediction scores from eight transcriptionally aware RFM models (GTEx-RFM; 1 indicating high confidence of editing and 0 indicating high confidence of no editing) providing a metric for dsRNA prediction confidence based on editability.

### Non-repetitive dsRNAs are enriched at chromosomal tips

Shifting focus, we next analyzed the genomic distribution of repetitive and non-repetitive dsRNAs to uncover patterns of enrichment across different regions of the genome. Repetitive dsRNAs were prevalent throughout the genome, averaging around 2 dsRNAs predicted per 1 kb of genomic sequence (1.7 dsRNAs/kb; **Figure 2A**) and were predicted in nearly every region of the genome, albeit some locations were more dense than others. Consistent with earlier reports inferring dsRNA based on editing sites,^6^ we predicted the highest density of repetitive dsRNA on chr19 which has a high abundance of repetitive elements.^36,37^

**Figure 2.**
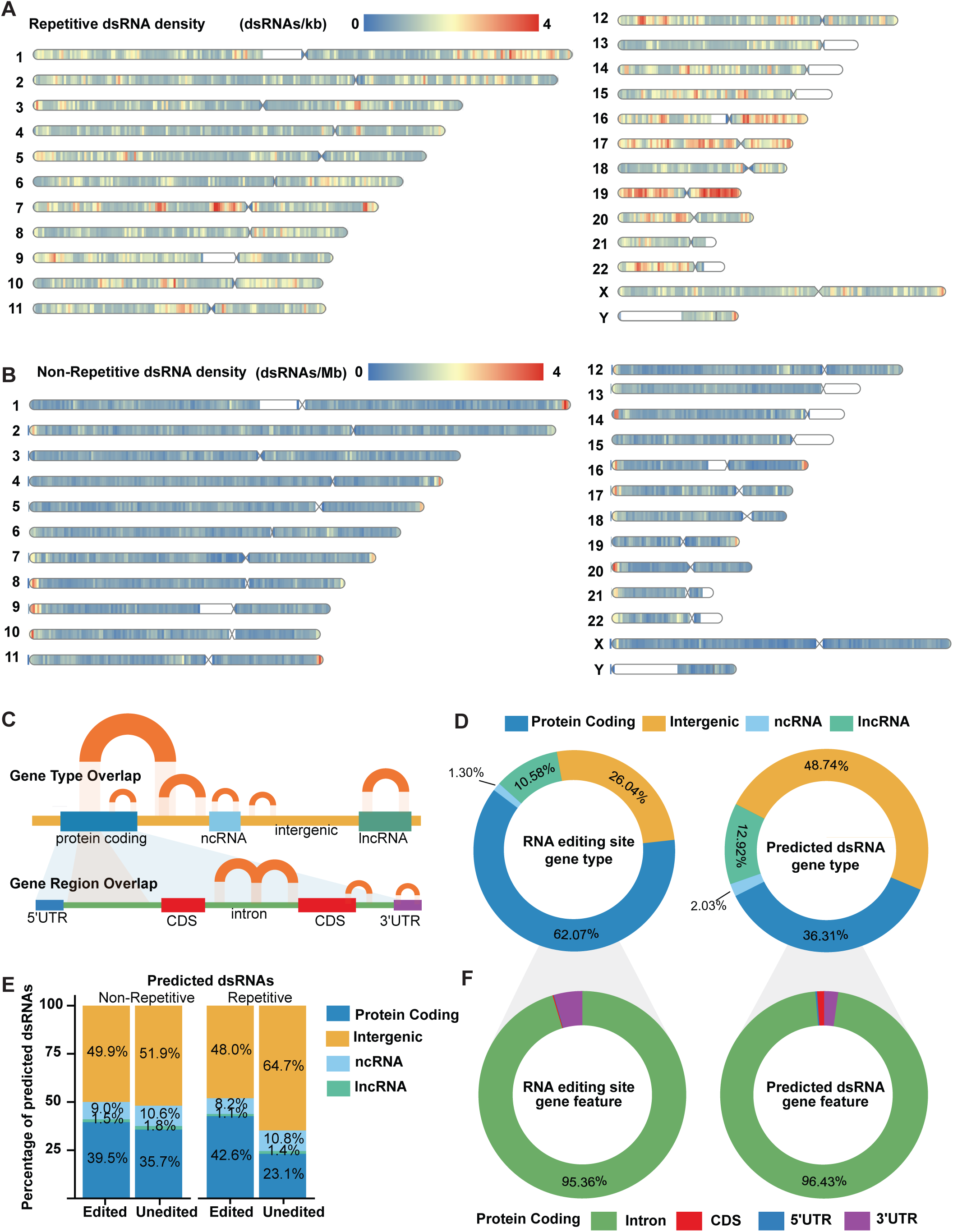
Genomic and genic distribution of predicted dsRNAs. **A.** Density, the number of predicted dsRNAs per kilobase (kb), for repetitive dsRNAs across all chromosomes, with a color gradient from blue (0) to red (4) dsRNAs per kb. **B.** Density of predicted non-repetitive dsRNAs per megabase (Mb) across all chromosomes, using color gradient as in A. **C.** Overlap of gene types and gene regions for predicted dsRNAs. Arcs represent overlap of dsRNAs with protein-coding genes, ncRNA, lncRNA, and intergenic regions. Overlap with gene regions includes 5’ UTRs, CDS, introns, and 3’ UTRs. A dsRNA was classified as part of a gene type/region if either arm overlapped with >95% of its nucleotides. **D.** Pie charts comparing proportion of predicted dsRNAs and RNA editing sites by gene type: protein-coding, intergenic, ncRNA, and lncRNA. **E.** Bar charts showing percentage of predicted dsRNAs by gene type (protein coding, intergenic, ncRNA, lncRNA) for non-repetitive and repetitive dsRNAs, split into edited and unedited categories. **F.** Pie charts illustrating distribution of predicted dsRNAs and RNA editing sites by gene feature: intron, CDS, 5’ UTR, and 3’ UTR.

Non-repetitive dsRNAs by comparison, were sparse, averaging around 2 dsRNAs per *mega*base of genomic sequence. For the most part these non-repetitive dsRNAs were spread out evenly throughout the genome, but interestingly, there was a consistent enrichment of non-repetitive dsRNA at chromosomal tips; 19/24 chromosomes had their highest non-repetitive dsRNA density within 3 million bps of their chromosomal tip (**Figure 2B**), it is unclear if this enrichment is connected to observations from *C. elegans* that found dsRNA enrichment at chromosomal tips as well.^38^

Prior studies show that RNA editing occurs most often in noncoding regions of pre-mRNAs (introns and UTRs) and less often in intergenic regions.^5–7,38,39^ However, it is unclear if reduced RNA editing in intergenic regions is due to lower levels of transcription, which makes editing detection a challenge, or if dsRNA forming sequences are depleted outside of gene regions. Importantly, dsRNAscan predicts dsRNA in an editing-neutral way, and thus offered the opportunity to answer this question. To this end, all dsRNAs were annotated based on their presence or absence in an annotated gene locus **(Figure 2C)**. While known RNA editing sites were mostly in gene loci, accounting for 74% of all editing sites, only 51% of our predicted dsRNAs were encompassed in genes (**Figure 2D**). Correspondingly, there was a large discrepancy in the percentage of editing sites and predicted dsRNA in protein coding genes (62.07% vs 36.31%), however, the proportion of editing sites and predicted dsRNAs in all other genes, were similar (12.92% and 10.58%, respectively; **Figure 2D**). The relative proportion of dsRNA overlapping repeats/editing sites was generally consistent across all categories with the exception of unedited repetitive dsRNAs which were more likely to be intergenic (**Figure 2E**). While intergenic repetitive dsRNA may be less likely to be edited, lower transcription levels in intergenic regions could also prevent the detection of RNA editing sites. Within protein coding gene loci, 96.43% of predicted dsRNAs and 95.36% of editing sites were found in introns (**Figure 2F**). In exons, RNA editing sites were rarely found, and dsRNAs were rarely predicted; when they were found, they were mostly in 3′UTRs, less so in the CDS, and most rarely in 5′UTRs (**Figure 2F**).

### Putative intramolecular MDA5 substrates are rare

We were particularly interested in dsRNAs predicted to be long and highly paired (>300 bps and >96% of nucleotides base paired) that might be substrates for MDA5, which cooperatively binds dsRNA to induce an interferon response; a process markedly favored by dsRNAs longer than 300 bp *in vitro*.^40,41^ Similar to previous computational studies,^42^ we found that long, highly paired dsRNAs were exceptionally rare in the genome. In our dataset, comprising a total of 5,134,754 dsRNA predictions, we found ∼1 million longer than 300 bps, 31,017 that were highly paired (>96%), but only 2,018 dsRNAs (0.00038%) that were *both* long and highly paired (**Figure S1A**).

While every chromosome harbored these long, highly paired dsRNA structures, chrX had the highest number (**Figure S1B**). For all chromosomes, we discovered that most of these formed between inverted LINE elements **(Table S5)**, which are most abundant on chrX.^43^ Most long, highly-paired dsRNAs were unedited (74.23%; **Figure S1B**), likely due to lack of transcription: 88% of long highly paired dsRNAs were intergenic, almost double the overall distribution of intergenic dsRNAs (48%; **Figure 2F**)—an overrepresentation that further suggested long, highly paired dsRNAs are less tolerated in gene loci.^42^ However, this led to the obvious question: are any long, highly paired dsRNAs expected to enter the cytoplasm?

To explore this question, we searched for dsRNAs in transcripts expected to be capped, polyadenylated, and transported to the cytoplasm like mRNAs or lncRNAs. Among the 2,018 long, highly paired dsRNAs, we only identified 58 that overlapped protein coding exons and 45 that overlapped lncRNA exons from Gencode V46. However, all but one of these were between an exon and an intron, which would result in the dsRNA being removed during splicing. Ultimately then, there was only a *single* long, highly paired dsRNA present in a 3′UTR: a ∼300 bp dsRNA embedded in the longest isoform of *ZNF426* (**Figure S2**), predicted to form between two highly similar inverted AluY elements, the youngest of the Alu elements which emerged around 10 million years ago. These results are consistent with the idea that highly immunogenic dsRNAs do not enter the cytoplasm under normal conditions.^44^

### Editing sites reveal hundreds of natural antisense dsRNAs

To date, there is little information about the frequency of intermolecular dsRNA formation, but increasingly, reports suggest this type of interaction may be biologically relevant. For example, recent studies suggest that immunogenic dsRNAs arise from intermolecular hybridization of natural antisense transcripts.^14,15^ dsRNAscan doesn’t predict intermolecular dsRNA, however, we reasoned that dsRNAscan could point to regions that were good candidates for intermolecular interactions. For example, editing sites that do not overlap predicted intramolecular dsRNAs indicate a dsRNA undetectable by our criteria, which could include dsRNA spanning >10,000 nts, short dsRNAs with <25 bps, intramolecular structures that form after splicing and bring distant arms together,^45^ RNA:DNA hybrids,^46^ or most notably for our analysis, intermolecular dsRNAs.^14,15,47^ We found that 93% of editing sites overlapped a predicted intramolecular dsRNA (**Figure 3A**). This designated the remaining 7% of editing sites, 1,137,706 in total to be further evaluated as candidates for intermolecular dsRNA formation.

**Figure 3.**
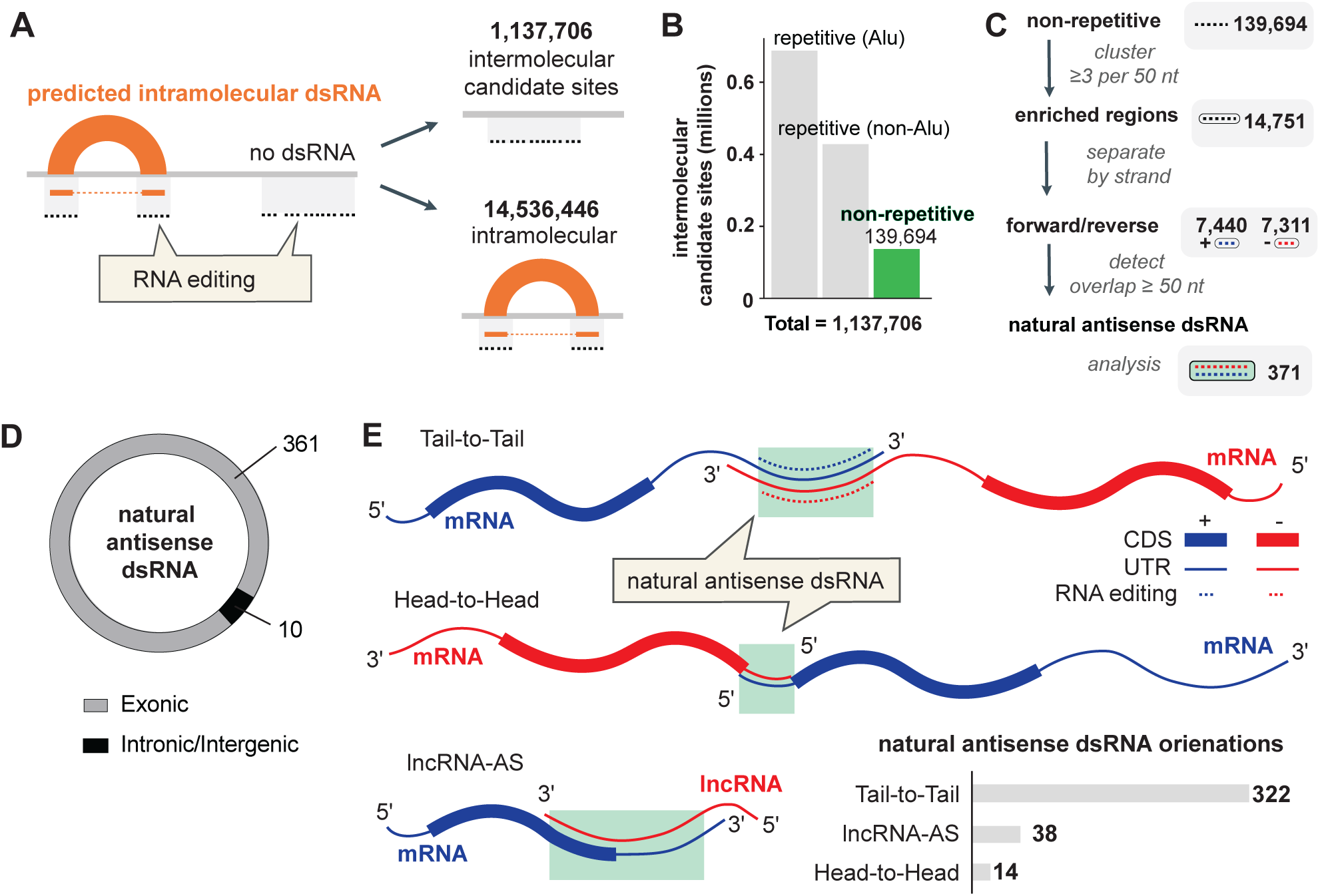
Prediction of intermolecular dsRNAs. **A.** Workflow for predicting intermolecular dsRNAs. Editing sites were intersected with dsRNA coordinates to find those that did not intersect with a predicted intramolecular dsRNA; these 1,137,706 sites were further analyzed as candidate intermolecular editing sites. **B.** Categorization of 1,137,706 candidate intermolecular dsRNAs as repetitive (Alu and non-Alu) and non-repetitive. **C.** Refining intermolecular candidates to find natural antisense dsRNAs. Editing sites within non-repetitive intermolecular candidate dsRNAs were used as input to find natural antisense dsRNA. These 139,694 editing sites were parsed into those that contained at least three editing sites per 50 nucleotides, defining 14,751 intermolecular candidate EERs, which were then split into separate strands and intersected with each other to find EERs that overlapped on both strands. EERs without complementary sequences in other genomic locations were presumed to form dsRNA with their natural antisense counterparts, resulting in 371 high-confidence natural antisense dsRNAs. **D.** Categorization of natural antisense dsRNAs based on their genomic location reveals a significant proportion in exonic regions. **E.** Schematic representation of different configurations of natural antisense dsRNAs: Tail-to-Tail, Head-to-Head, and lncRNA-AS. Orientation numbers for bar graph (Bottom Right) were approximated based on genomic overlaps of each dsRNA, such as their presence within annotated 3’UTRs, 5’UTRs, or lncRNAs.

Further analysis revealed that 77% of these intermolecular candidate editing sites were in repetitive elements (**Figure 3B**); non-repetitive sites are only 2.75% of all editing sites yet contributed 12.47% of the intermolecular candidates (**Figure S3**). When considering *repetitive* intermolecular candidate sites, their existence was anticipated and intermolecular hybridization of inverted repetitive elements is a proposed mechanism of how DNA-demethylating agents induce dsRNA formation^48,49^; the sheer abundance of repetitive elements in the transcriptome hinders attempts to predict which ones hybridize to form intermolecular structures (**see Discussion**). Therefore, to maximize the likelihood of identifying *bona fide* intermolecular dsRNAs, we focused on non-repetitive sites as the first step in identifying natural antisense dsRNAs (**Figure 3C**).

First, we clustered the 139,694 non-repetitive, intermolecular candidate sites into groups; non-repetitive intermolecular candidate sites were considered an editing enriched *region* (EER) if it had ≥3 editing sites per 50 nt (**Figure 3C**). Grouping editing sites into regions is a well-established method to facilitate identification of long dsRNAs.^5,6,39,50^ This resulted in 14,751 intermolecular candidate EERs. To determine if these regions were forming dsRNA due to natural antisense pairing, which occurs when the same region of the genome is transcribed in both directions, we used BLAST to check for the presence of complementary sequences elsewhere in the genome. We focused on regions that were at least 25 bps long with over 90% sequence identity. We found that only 1,030 (7%) of the intermolecular candidate EERs had complementary sequences in other parts of the genome, typically found on different chromosomes or in distant intrachromosomal regions (>10,000 nt away), usually forming between low complexity or repetitive gene loci such as ZNF^51^ genes (**Table S6**). These regions are therefore capable of forming dsRNA through interactions with these distant sequences rather than with only their natural antisense transcripts. The remaining regions, which did not show significant BLAST hits, were assumed to form dsRNA with their natural antisense partners. To increase confidence in this hypothesis, we next focused on regions with editing sites on both strands – a trait that can be indicative of intermolecular dsRNA formation.^52^ This criteria resulted in 371 high confidence natural antisense dsRNAs (**Figure 3C**), almost all of which (97%) were found in gene regions, and consistent with their proposed regulatory roles, almost 80% were in 3′UTRs (**Figure 3D**). The enrichment of 3′UTRs in these results further established these could represent *bona fide* regulatory intermolecular interactions.^53^

To better understand the spatial organization of natural antisense dsRNA formation, we categorized the 371 high-confidence natural antisense dsRNAs into three structural classes based on how their transcripts overlapped: tail-to-tail, head-to-head, and lncRNA-antisense configurations (**Figure 3E**). In the tail-to-tail arrangement, the 3′ ends of two transcripts overlap, forming dsRNA between their complementary regions, which was the most prevalent orientation. In contrast, the head-to-head configuration, where the 5′ ends of two transcripts overlap, was much rarer. The lncRNA-antisense configuration, involving a long noncoding RNA overlapping a protein-coding gene in an antisense manner was counted 47 times.

### Conserved dsRNAs are less structured, rarely edited, and enriched in neurons

While the millions of ADAR editing sites in noncoding sequences indicate they are far more abundant, there are also hundreds of conserved editing sites in the human genome that recode proteins.^54–57^ One of the most recognized is the Q/R editing site in *GRIA2*’s coding sequence (CDS)^58^, which must be edited for proper glutamate receptor function. In the absence of editing, GRIA2 is non-functional resulting in embryonic lethality.^59^ Such crucial recoding editing sites are under immense evolutionary pressure, resulting in conservation of their underlying dsRNA secondary structures, finely tuned for ADAR editing.^60^ However, excluding recoding sites, it is unclear if any other conserved dsRNAs exist. To address this, we again took advantage of dsRNAscan’s editing agnostic approach.

All dsRNA arms were overlaid with two sets of phastCons scores (**Figure 4A)**, which estimate the conservation of each nucleotide across 100 vertebrates and 17 primates (phastCons100 and phastCons17); high scores (closer to 1) suggest evolutionary pressure to maintain sequence, while low scores (closer to 0) suggest variability and lack of evolutionary constraint.^61^ For each arm the average phastCons score was calculated: over 99% of predicted dsRNAs had phastCons scores below 0.5 and only 1,502 dsRNAs had a score ≥0.9 in 100 vertebrates and 17 primates (**Table S7**).

**Figure 4.**
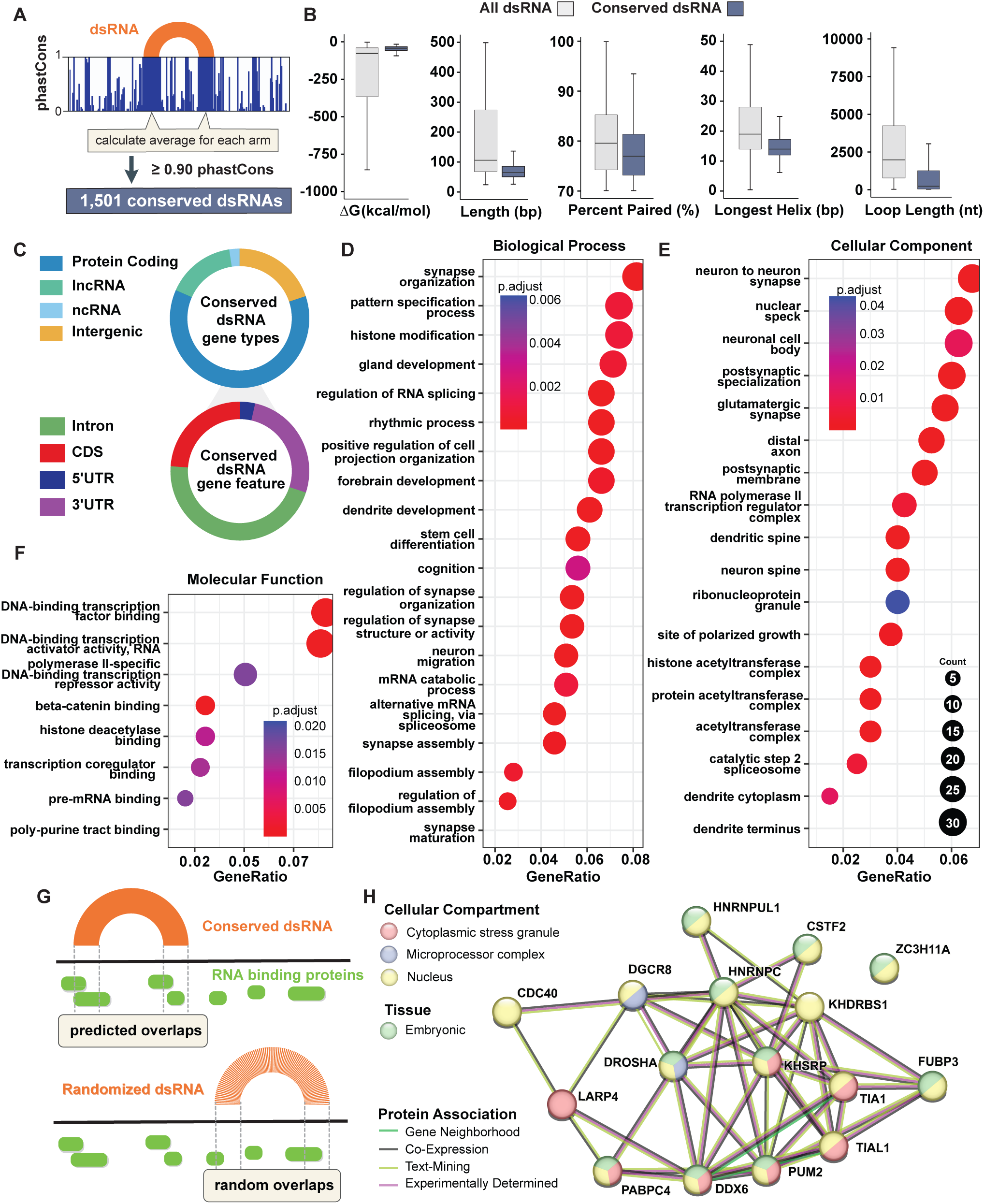
Structural and functional analysis of conserved dsRNAs. **A.** Schematic representation of assigning conservation data to dsRNA. Each arm of a dsRNA consists of coordinates which we used to calculate the average phastCons score for that coordinate range. We defined a dsRNA as conserved if both arms have an average phastCons > 0.9 for both phastCons17 and phastCons100, which estimate conservation of the sequence across 17 primates and 100 vertebrates, respectively. These criteria revealed 1,501 conserved dsRNA. **B.** Structural properties of conserved dsRNAs. Box and whisker plots depicting distribution of structural characteristics of conserved dsRNA (n = 1,501) vs. all dsRNA (n = 5,134,754). Boxes represent interquartile range (IQR), with horizontal line indicating median value. Whiskers extend to minimum and maximum values within 1.5 times the IQR. Outliers were removed for clarity. **C.** Classification of conserved dsRNAs by gene type and feature. Conserved dsRNAs were tested for overlap with gene types: 927 overlapped protein coding genes, 240 overlapped lncRNAs, 36 overlapped ncRNAs, and 391 were intergenic. Within the 927 protein coding genes the majority overlapped introns (466) but were also found to overlap coding sequences (CDS; 241), 3’UTRs (267), 5’UTRs (36). If a dsRNA overlapped multiple gene types or features, it was counted in each category, resulting in totals that may exceed the number of unique dsRNAs. **D.** GO Biological Process enrichment analysis. Dot plot displays enriched biological processes for conserved dsRNA-containing genes. Each dot represents a specific GO term; size correlates with number of input genes in each category (see legend on right side of Panel E) with GeneRatio on x-axis reflecting fraction of input genes associated with that term. Larger dots are terms with more associated genes, and color gradient from blue to red is adjusted p-value (p.adjust) after Bonferroni correction, with darker red dots indicating more significant terms. For details see Methods. **E.** GO Cellular Component enrichment analysis. Dot plot presents cellular components associated with conserved dsRNA genes. **F.** GO Molecular Function enrichment analysis. Plot shows molecular functions most enriched among conserved dsRNA-containing genes. **G.** Comparison of predicted vs. randomized dsRNA RBP binding sites. Schematic representation showing predicted RBP overlaps on conserved dsRNAs versus randomized controls. **H.** RBPs enriched on conserved dsRNAs. STRING network analysis of RBPs that show significant enrichment (p < 0.005) on conserved dsRNAs. Nodes represent specific RBPs, colored according to gene ontology/tissue enrichment. Connecting line colors represent interaction confidence based on protein-protein associations across various datasets.

Structurally, conserved dsRNAs represented the least stable dsRNA category of our study, yielding just -42 kcal/mol free energy prediction on average (**Figure 4B**; **Table 2**). Compared to all predicted dsRNAs, conserved dsRNAs were shorter, had a lower percentage of paired nucleotides, shorter stretches of contiguous bps, and shorter loop lengths (**Figure 4B**; **Table 2**). Accordingly, conserved dsRNAs were rarely edited; only 29 overlapped an editing site on both arms and only 81 had at least one editing site on either arm (**Table S7**).

**Table 2.**
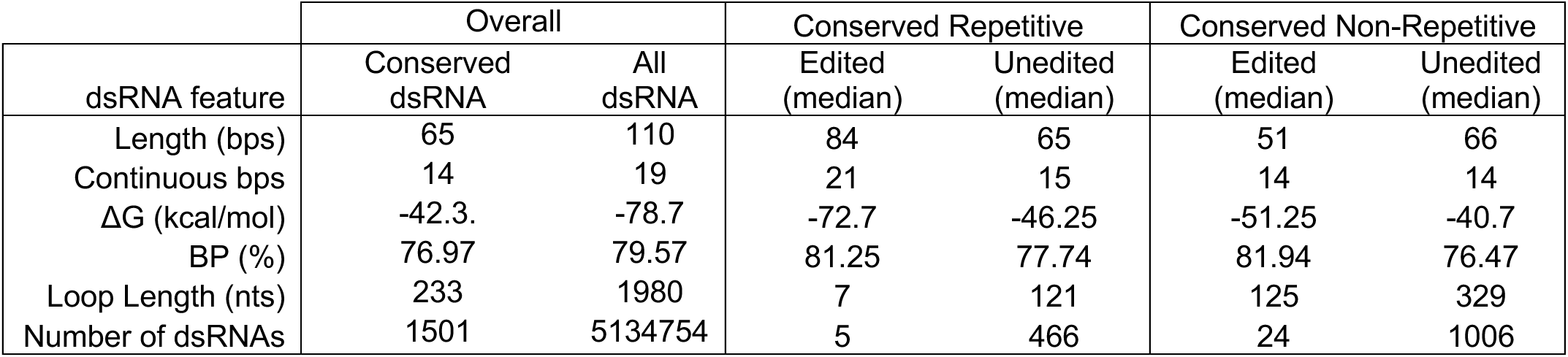
Summary of structural characteristics for conserved dsRNAs. This table contains structural metrics for conserved dsRNAs that are edited and unedited dsRNAs across repetitive and non-repetitive regions of the human genome compared to overall dsRNA characteristics. Median values for key features, including dsRNA length, contiguous base pairs (bps), free energy of formation (ΔG), base-pair percentage (BP%), and loop length are reported for both repetitive and non-repetitive dsRNAs.

Conserved dsRNAs were enriched in gene regions, with 927 conserved dsRNAs overlapping 513 genes, including 416 protein coding and 97 lncRNAs (**Figure 4C**) as well as 45 primary miRNA stems (**Table S8**). GO enrichment analysis of genes with conserved dsRNAs revealed significant involvement in biological processes such as synapse organization and neuron projection regulation (**Figure 4D**). Conserved dsRNA genes were also enriched in cellular components like postsynaptic membranes, dendritic spines, and neuronal cell bodies (**Figure 4E**), suggesting a specific role in synaptic organization and dendritic development. Furthermore, molecular function analysis showed strong associations with transcriptional regulation, including binding of DNA-binding transcription factors and transcription co-regulators (**Figure 4F**). Additionally, we tested if genes with conserved dsRNAs were tissue specific^62^, finding that out of the 35 tissues catalogued in the Human Protein Atlas there was enrichment in only the cerebral cortex (62 genes; **Table S9**).

Given these strong correlations with neuronal function, we next shifted our focus to individual dsRNA to explore their conservation and structural properties in more detail. While all structures were potentially interesting due to their conservation alone, we prioritized those with the highest confidence of formation. Inclusion of editing sites provides support for the existence of a dsRNA, but for the 1,420 dsRNAs without editing sites, we used our machine learning, transcription-aware model to rank their likelihood of being edited. We predicted 87 dsRNAs with a greater than 50% chance of editing (i.e., a model score >0.50; **Table S7**). The highest scoring dsRNA (average GTEx-RFM score of 0.97) was predicted to form between an intron and an alternative 3′UTR isoform of SON (**Figure S4**), a protein with a dsRNA binding domain (dsRBD) involved in neurodevelopment and alternative splicing. This dsRNA was 180 bp long, composed of three helices interrupted by mismatches and bulges, and a long “loop” region (**Figure S5**). Consistent with machine learning predictions, the first helix contained three adenosines that were highly edited in brain tissue as reported in REDIportal (**Figure S6**), and all helices showed strong conservation across 176 vertebrate species, with helix three displaying statistically significant covariation (**Figure S7).** Identifying novel RNA structures through covariation analysis is particularly challenging in mammals,^63^ but when observed, it provides compelling evidence that the dsRNA is functionally significant, as seen with the SON dsRNA.^64^

One of the most common ways RNA structures impart function is by presenting binding sites to specific RNA-binding proteins (RBPs). Thus, we were interested if any RBPs were predicted to bind to these conserved dsRNAs. To explore this, we analyzed eCLIP data, which maps binding locations of 150 RBPs across the transcriptomes of two cell lines.^65,66^ We determined overlap between RBP binding sites defined by eCLIP data, and conserved dsRNA regions as well as randomized control regions (**Figure 4G**). By applying statistical tests (see Methods), we determined whether RBP binding sites intersected dsRNAs more than expected by chance. We identified **17** RBPs that showed significant enrichment for binding to dsRNA regions compared to random controls (**Figure 4H**). Notably, the RBPs with the most significant enrichment were TIA1 and KHSRP, proteins involved in neuronal development^67–69^ and neurodegenerative disease,^70,71^ with both predicted to bind the conserved dsRNA in SON as well.

## DISCUSSION

In this study, we developed and utilized dsRNAscan to generate a comprehensive map of dsRNAs encoded in the human genome. Predictions were made independent of RNA editing to overcome several limits of RNA-seq-based approaches, such as the need for transcription and ADAR enzyme activity. This allowed for the prediction of over five million dsRNAs that are both edited and unedited, and represent a range of structural characteristics (**Figure 1**). These diverse dsRNA predictions provide an important resource that will drive discoveries across many fields. For example, we observed an enrichment of non-repetitive dsRNAs near chromosomal tips (**Figure 2B; Table S2**), setting the stage for insights into how RNA impacts genome architecture.^72^ We catalogued hundreds of conserved dsRNAs enriched in genes associated with neurodevelopment, underscoring the already intriguing role of dsRNA in neurology **(Table S7)** and highlighting specific dsRNA structures that may have unique functional roles. While the dsRNAs that trigger an immune response in mammalian cells deficient for ADARp150 are unknown, we identified a subset of dsRNAs that are good candidates for the elusive immunogenic dsRNAs (**Table S5**), paving the way for a mechanistic understanding of aberrant immune responses and autoimmune disease. These findings emphasize how dsRNAscan and its results for the human genome will spur new research towards mechanistic insight.

### Utilizing dsRNA predictions in research and therapeutics

The dsRNAscan results for the human genome (www.dsrna.chpc.utah.edu) offers researchers a convenient way to explore the presence and predicted base pairing of dsRNA structures across the human genome. We provide tools to filter and download dsRNAs, allowing researchers to focus on dsRNAs in their genes of interest. As described in our user guide (**Document S1)**, users can quickly home in on interesting structures using the IGV genome browser or download dsRNAs in bulk as spreadsheet files. Integrating dsRNA predictions with genomic datasets will reveal dsRNAs that can be tested for roles in key processes like alternative splicing or transcript stability, even for unedited dsRNAs. For example, overlaying dsRNA with phastCons scores revealed hundreds of unedited but highly conserved dsRNAs that could function as gene regulatory features during neurodevelopment.

Additionally, our analysis of RBP binding sites uses scripts (shared at https://github.com/Bass-Lab/) that can be refactored to determine if dsRNAs associate with other genomic datasets, such as the locations of circRNA junctions,^73^ polyadenylation sites,^74–78^ and intron-exon junctions.^79^ Similarly, correlating dsRNA predictions to specific regulatory outcomes suspected to involve dsRNA such as nonsense-mediated decay,^80^ Staufen-mediated decay,^81^ intron retention,^82,83^ splicing efficiency,^84^ or alternative splicing,^85^ may reveal a layer of regulation via dsRNA formation. The ability to map precise locations and base-pairing patterns of dsRNAs allows researchers to easily check if dsRNA formation might influence post-transcriptional regulation by occluding or exposing splice sites, miRNA binding sites, or polyadenylation signals.

Beyond exploring how dsRNAs impact RNA biology, dsRNAscan will provide new information in the design of therapeutic strategies, for example those focused on autoimmune disorders and RNA-based therapies. By examining structural characteristics predicted by dsRNAscan, researchers will identify the location of potentially immunogenic dsRNAs promoting insights into the molecular triggers of autoimmune disorders and cancer. For instance, are there expressed intergenic loci that correlate with specific cancer or immune diseases? Being able to quickly determine if such regions can form dsRNA will focus efforts to understand pathogenicity. Likewise, understanding which genomic regions can form immunogenic dsRNAs opens new therapeutic avenues. For example, CRISPRa^86^ could be employed to increase transcription of immunogenic dsRNAs, potentially activating the immune system as part of a viral mimicry therapeutic strategy.^82,87–91^

### Insights into protein binding

Separating predicted dsRNAs into edited/unedited categories provided insights into the preferred human ADAR substrate (**Table 1**). Since most repetitive and edited dsRNAs are formed by Alus, the structural characteristics of this category mostly align with Alu sequence characteristics (i.e. average length of repetitive, edited dsRNA and Alus is both ∼300 bp). On the other hand, when comparing edited vs unedited non-repetitive dsRNA, we observed differences that point to ADAR’s preferred dsRNA substrate: editing occurred in longer substrates (**Figure 1E**) with smaller loops (**Figure 1I),** and within these, a contiguous stretch of longer than 30 base pairs was preferred (**Figure 1F**); unedited dsRNAs were shorter (**Figure 1E)**, with longer loops (**Figure 1I**), with a median contiguous base pair stretch of only 14 bp (**Figure 1F**). This is consistent with the current understanding that most editing is found in longer dsRNAs, while ADAR substrates with less than 17 bps are exceptional.^92,93^ It is important to emphasize that unedited dsRNA structures may also have functional roles. Indeed, some of our predicted structures *are* functional, such as the conserved dsRNAs that comprised known primary miRNA structures (**Table S8**). These structures are sometimes edited, but are often unstructured enough to inhibit ADAR editing while still allowing for dsRNA binding by DROSHA and DGCR8.^94–97^

This highlights that smaller or otherwise “suboptimal” dsRNAs—while not edited by ADARs—may still be recognized by other RBPs. RBPs interact with their RNA targets through a combination of sequence motifs and structural elements^98^; RBPs use modular arrangements of domains like RRMs (RNA recognition motifs), zinc fingers and KH domains to bind single-stranded regions, while domains like the dsRBD engage with dsRNA helices.^99^ Almost all conserved dsRNAs we observed had a mix of paired and unpaired regions of RNA which might offer uniquely spaced motifs for specific RBPs. There are reportedly hundreds of dsRNA binding proteins (dsRBPs) with novel modes of interaction, like the ARM domain^100^ and over 1500 RBPs with unknown mechanisms of binding—with genome-wide efforts underway to reveal their binding preferences.^65,98,101,102^ This allows for the possibility that some of these unedited dsRNAs may provide specialized structural and sequence motifs for individual RBPs. Our dataset provides a map of these structures ensuring they can be easily considered in future studies. This will help broaden our view of dsRNA and its potential role in cellular regulation beyond editing and immunogenicity.

### Is there a function for dsRNAs in neurogenesis?

Neurons have demonstrated a unique relationship with dsRNA: neurons have the highest level of RNA editing,^3,4^ the pre-mRNA of neuronal receptors, such as GRIA2, encode specialized dsRNA structures that promote precise ADAR binding and editing^31,60^; neurons are the only cells that express ADAR3, a non-catalytic member of the ADAR family that binds dsRNA and inhibits RNA editing,^103^ modulating immunity,^104^ and contributing to oncogenesis^105^; it was also recently discovered that neurons transcribe longer 3′UTRs in order to present immunostimulatory dsRNAs to dsRNA sensors in the brain,^106^ thereby priming the cellular immune system and enhancing antiviral defense.^107^ In this context, we believe our finding of hundreds of highly conserved dsRNAs enriched in the cerebral cortex and in neurologically related genes is particularly interesting. The conservation of these dsRNAs across vertebrates and primates suggests they could have specific functions that pressure their formation. Further analysis of these regions could prove insightful and as an example, we presented an analysis of one conserved structure formed in the SON pre-mRNA (**Figures S4-S6**).

The conserved structure in SON presents several intriguing characteristics worth discussing. Its editing in the brain (**Figure S6**) provides strong evidence of its formation *in vivo,* and the presence of a significantly covarying base pair (**Figure S5**) increase the likelihood it serves some functional role. Intriguingly, this dsRNA forms between an intron and the 3′UTR of a short SON isoform (**Figure S4**). In the longest isoforms of SON, this conserved dsRNA would be spliced out completely, indicating that its formation is uniquely regulated in the short isoform, where it becomes part of the 3′UTR. This short isoform, notably, does not encode the dsRBD of SON, allowing for the possibility of autoregulation where the full-length *SON*, or other RBPs, can bind the dsRNA and modulate the splicing of the short isoform. In this model, the dsRNA would serve as a regulatory switch, modulating the recruitment of splicing factors in a context-dependent manner.

### What are the most likely endogenous substrates activating MDA5?

RNA editing by ADARs plays a critical role in preventing endogenous dsRNAs from activating MDA5, which recognizes long, highly paired dsRNAs—structures that resemble viral replication intermediates. Small structural disruptions, such as bulges or mismatches, which are common in dsRNAs formed by virtually all inverted Alu elements, typically impair MDA5 activation.^98^ Additionally, only a small number of ADAR editing sites in a given dsRNA are required to prevent autoimmune responses in this way.^15^ Consistently, we identified only a few thousand long, highly paired dsRNAs (>300 bp and >96% paired) across the genome (**Table S5**). However, according to current genome annotations, these dsRNAs were almost non-existent in mature mRNAs, consistent with the idea that they are largely selected against.^42^ In fact, the only long highly paired dsRNA located in a 3′UTR (of ZNF426), was formed by Alu elements from the *youngest* class, Y, which first appeared around 10 million years ago.

In addition to these intramolecular dsRNAs, we identified hundreds of natural antisense dsRNAs (**Table S6**). These dsRNAs are predicted to form between sense and antisense transcripts, and have recently been suggested to be some of the most potent immunogenic dsRNAs.^14^ The longest natural antisense region we defined was almost 900 bp long, however many of these regions had neighboring natural antisense regions within a few hundred nucleotides, suggesting natural antisense dsRNAs could grow longer than 1000 bp, a length shown to activate MDA5 more robustly *in vivo.*^108^ In many ways these structures mimic viral dsRNAs more closely than inverted Alus, making them reasonable candidates for MDA5 activation. However, the timing of their formation—whether they are edited in the nucleus or hybridize after nuclear export—remains unclear, but this presents an important question for further research. Knowledge of the location and expression of these natural antisense dsRNAs (**Table S6**) could lead to new strategies for testing hypotheses, combatting immune activation in diseases like Aicardi-Goutières syndrome, or designing strategies to induce immune activation to fight cancer.

### Considerations and limitations

To maintain computational feasibility, einverted reports the highest scoring, longest, inverted partner for a given sequence. Reporting all suboptimal, overlapping, and competing inverted repeats would exponentially increase computational costs. Therefore, in windows with dozens of inverted repeats, suboptimal and competing intramolecular dsRNAs are omitted in favor of the highest scoring partners. Similarly, since repetitive elements like Alus are so abundant—found in almost all genes—they are capable of making intermolecular contacts between any of their inverted partners being transcribed across the genome. Because of these complications we focused our analysis of natural antisense dsRNAs on non-repetitive regions. However, it is likely that intermolecular dsRNAs are even more prevalent between repetitive regions, such as Alu elements, where broad sequence similarity facilitates pairing. While many of these Alu are presumed to be edited after forming intramolecular dsRNA, our analysis uncovered thousands of editing sites in repetitive elements that did not correlate with intramolecular dsRNA formation. Trying to determine if these are formed intermolecularly or intramolecularly will require newer techniques like long-read sequencing methods.^109^

In order to wholly map endogenously coded dsRNA across the human genome, we assumed that any given 10,000 nt stretch of DNA could be transcribed into RNA. Because of this, we must consider that some dsRNA would only form if the entire region is transcribed. For example, four of our conserved dsRNAs were predicted to form between inverted tRNA genes; some of these examples were in introns which are transcribed and would allow for dsRNA formation, but others were in otherwise silent regions of the genome. Therefore, it is important to consider the full annotation context of a predicted dsRNA when considering the likelihood of its formation.

Our default dsRNAscan settings defined dsRNA as having a minimum length of 25 bps. This length cutoff was intended to avoid excessive predictions and indeed, 99% of predicted dsRNA were between 30 and 1000 bp. However, shorter dsRNAs and/or dsRNAs with minimal disruptions are still recognized by dsRBDs, as isolated mismatches can engage in non-Watson-Crick pairing or offer sequence specific interactions.^110^ Because of this, we believe another scan focused on finding “suboptimal” dsRNAs unlikely to be immunogenic or ADAR substrates, could be fruitful. For example, the properties of conserved dsRNA were relatively unstable and rarely edited (**Table 2**). This could prove to be a hallmark of dsRNA with functional roles, whereby they adopt enough structure to serve as a dsRNA substrates but also have enough disruptions—via internal loops, mismatches or bulges—that they are ignored by the immune sensors.

## Acknowledgements

We thank members of the Bass Lab for helpful discussions and feedback. We thank Supraja Ranganathan for computational help while preparing machine learning training data. We would also like to thank Enrique Arce-Larreta for help with machine learning strategy. The support and resources from the Center for High Performance Computing at the University of Utah are gratefully acknowledged. This work was supported by funding to R.J.A. from the National Institute of General Medical Sciences (F32GM153132) and to B.L.B from National Institute of General Medical Sciences (R35GM141262) of the National Institutes of Health. B.L.B. is a Jon M. Huntsman Presidential Endowed Chair.

## Author Contributions

R.J.A designed and performed the experiments, analyzed the data, and drafted the manuscript.

B.L.B supervised the research, provided guidance throughout the study, and contributed to the writing and revision of the manuscript.

## Methods

### dsRNAscan pipeline

The dsRNAscan pipeline was implemented using Python and designed to predict intramolecular dsRNA structures across the entire genomic sequence. The dsRNAscan pipeline utilizes the *einverted* algorithm, a core component of the EMBOSS package^17^ that specializes in detection of inverted repeats that form base paired regions of a dsRNA helix. The dynamic programming algorithm factors in both matches, which increase the score and represent base pairs, and mismatches or gaps, which incur penalties and represent bulges, an allowance that allows for dsRNAs with bulges and internal loops scattered throughout.^111^ A critical step in designing dsRNAscan was configuring and picking an optimal score for identifying potential dsRNA structures.

The target we chose for these configurations was derived from the well-characterized dsRNA structure at the GRIA2 (Glur2/GlurB) Q/R site.^112^ This site, encompassing a length of 30 base pairs, contains 8 GU pairs and exhibits a 93% match rate, inclusive of 2 mismatches. To ensure the accurate identification of similar structures, the einverted algorithm was modified to enable recognition and scoring of GU pairs as matches. While the default parameters of einverted — match and mismatch scores of +3 and -4, respectively, and a gap penalty of 12 — yielded a score of +20 for the GRIA2 Q/R site, our updated parameters scored this site +76. This heightened sensitivity, however, resulted in a dramatic increase in the number of dsRNAs identified. To maintain a stringent detection criterion and avoid excessive false positives, we set the default score threshold to 75.

When configuring default dsRNAscan parameters, selection of appropriate window and step size was important for balancing computational feasibility and accurate assumptions about RNA structure formation. We settled on a window size of 10,000 nt, which is both computationally tractable and sufficiently wide to encompass the majority of known intramolecular RNA-RNA interactions.^32^ This size ensures the capture of most RNA structures, and implicitly favors the detection of kinetically favorable dsRNA formed locally. Complementing this, a step size of 150 nt was utilized in order to increase prediction coverage while remaining computationally manageable.

Next, the predicted dsRNA structures are refined and analyzed using RNAduplex^11,18,19^ (Version 2.5.1), a program that predicts the optimal thermodynamics of hybridization between two RNA strands. This adds a “check” to dsRNA predictions and results in a valuable thermodynamic quantification for all predicted dsRNAs. Duplicate predictions were removed, and all predictions that completely overlapped each other on *both* arms were grouped together, with only the longest being reported.

### Machine learning

To assign confidence scores to dsRNA predictions, we leveraged machine learning models trained on over 5 million dsRNA predictions from dsRNAscan for the human genome. We aimed to classify dsRNAs based on key features like RNA editing status, nucleotide composition, base-pair counts, loop lengths, energy metrics, and conservation scores. The dataset was first preprocessed by calculating the average number of edits per 50 nucleotides to categorize dsRNAs as edited or unedited. To mitigate class imbalance, we applied SMOTE, ensuring equal representation of both classes. We also converted all numerical data to float32 format to optimize memory usage and computational efficiency. The dataset was then divided into training (80%) and test (20%) sets. Two feature sets were designed: a “No-GTEx” model, focusing on core dsRNA features like base-pair counts and trinucleotide compositions, and a “GTEx” model, which incorporated tissue-specific expression data alongside core dsRNA features to explore relationships between dsRNA formation and tissue-specific factors. We implemented two decision forest models—Random Forest and Gradient Boosted Trees— using the TensorFlow Decision Forests library.^33,34^

Both models were constructed with 1000 trees and were trained with varying tree depths from 2 to 8, optimizing for classification performance. Early stopping was employed to prevent overfitting, halting training if the validation loss did not improve for five consecutive epochs. Model performance was evaluated across depths using metrics like accuracy, log loss, precision, recall, and F1 score, which allowed us to select the optimal tree depths.

After model training and evaluation, we applied the models to the full dsRNAscan dataset, assigning confidence scores to each dsRNA prediction based on their structural and sequence characteristics. These confidence scores classified dsRNAs as either edited or unedited and were integrated into the dsRNAscan datasets.

### Gene Ontology (GO) Enrichment Analysis

GO enrichment analysis was performed to identify biological processes, cellular components, and molecular functions associated with conserved dsRNA-containing genes. The analysis was conducted using the clusterProfiler package in R, with gene IDs converted from Ensembl to Entrez using the org.Hs.eg.db annotation package (v3.14.0). Genes with conserved dsRNAs were input into the enrichGO() function to assess their enrichment within GO terms from the “Biological Process” (BP), “Cellular Component” (CC), and “Molecular Function” (MF) categories. Bonferroni correction was applied to account for multiple hypothesis testing, ensuring that only terms with a q-value below 0.05 were considered significant. Redundant GO terms were minimized using the simplify() function, which clusters semantically similar terms based on a cutoff of 0.7 and selects the term with the smallest adjusted p-value. The enrichment results were visualized using dot plots. All analyses were performed using the latest version of clusterProfiler and associated Bioconductor packages, with required package updates managed via BiocManager. Data visualization was performed with ggplot2 and enrichplot, with figures exported in SVG format for high-resolution display.

### Tissue Enrichment

Tissue enrichment of genes containing conserved dsRNAs was performed using the TissueEnrich package^62^ from its webserver on September 13, 2024. All gene symbols from **Table S7** were tested and the dataset was set to Human Protein Atlas.^113^

### Conserved dsRNA

In order to build consensus structure alignments, we blasted conserved dsRNA regions, including the loop region. We used blastn and searched all vertebrate genomes from the genomic reference sequence database. All hits with >90% query coverage were aligned using MAFFT. The resulting alignment was run through R-scape to predict a conserved secondary structure with the “--cacofold” and “--decoding” options. Importantly, we provided no secondary structure to R-scape, instead inferring the most likely structure based on alignment statistics when possible.^114^ The resulting Stockholm file was then analyzed with R2R^115^ to build the most informative consensus structure. In order to focus on the conservation of the human structure, we removed gaps from the human alignment sequences.

### Data acquisition

In our analysis, we utilized the GENCODE V46 annotations for gene annotations.^116–118^ RNA editing coordinates were obtained from REDIportal, which aggregates editing sites derived from over 9,642 RNA-seq samples.^3,4^ Tissue-specific expression data was retrieved from the GTEx database to incorporate into transcription-aware models. Conservation scores were calculated using phastCons data from the UCSC Genome Browser, representing sequence conservation across 100 vertebrates and 17 primates.^61^ Tissue enrichment of genes with conserved dsRNAs was performed using the TissueEnrich webserver.^62^ Finally, eCLIP data from ENCODE was analyzed to identify RBP binding overlaps with predicted dsRNA regions. ^65^

### Data analysis

R and python scripts were used to analyze data and generate figures and can be found at https://github.com/Bass-Lab/dsRNAscan.

---

**Figure S1.**
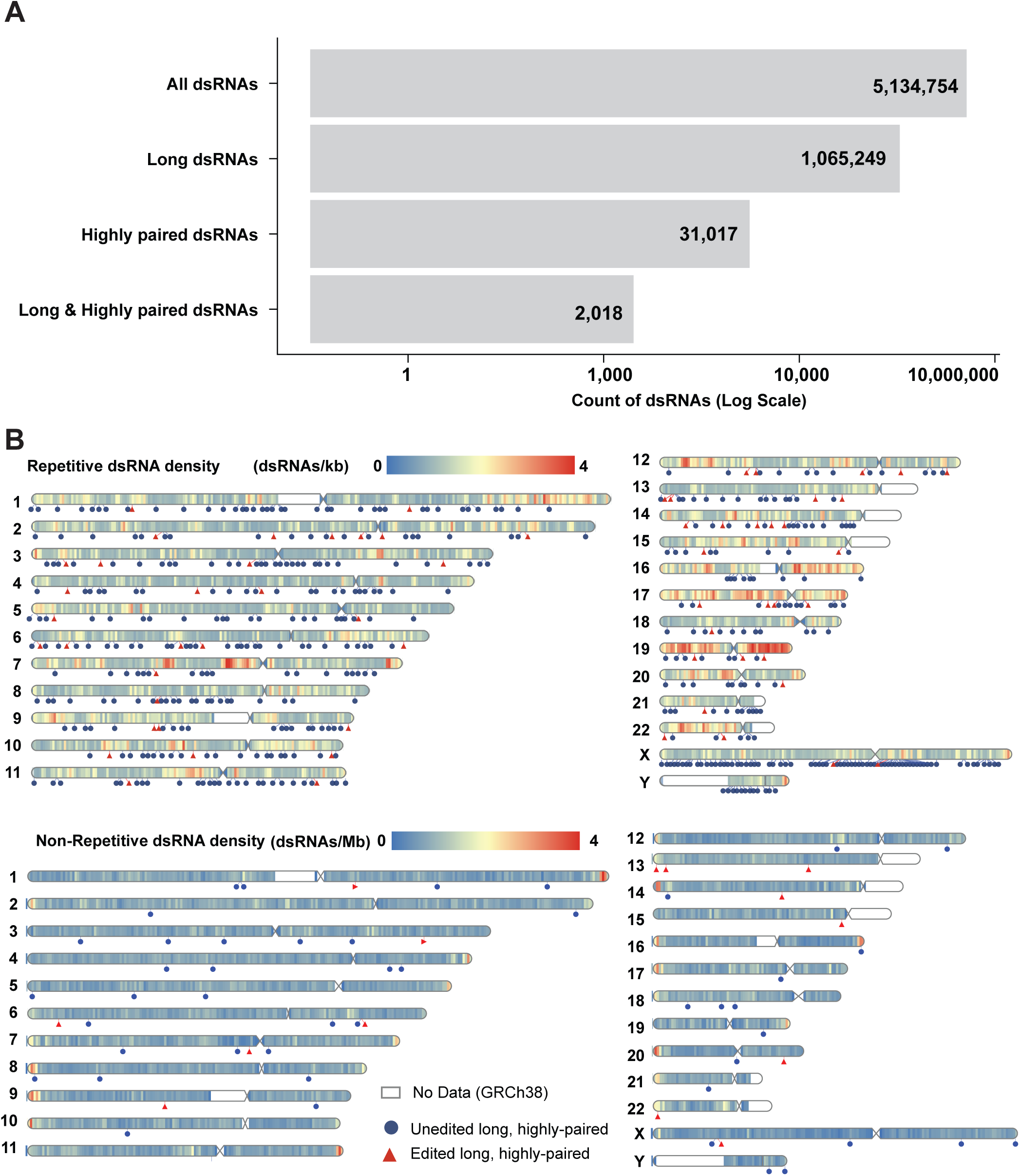
Genomic distribution of long dsRNAs. **A.** Bar graph showing the relative quantity of all dsRNAs vs long (>300 bps), highly-paired (>96% nts base paired), and long, highly-paired (>300 bps & >96% percent paired). **B.** Density, the number of predicted dsRNAs per kilobase (kb) for repetitive and per megabase (Mb) for non-repetitive dsRNAs across all chromosomes, with a color gradient from blue (0) to red (4). The location of long, highly-paired dsRNA (>300 bps and >96% bases paired) are marked on each chromosome by blue circles (unedited) and red triangles (edited).

**Figure S2.**
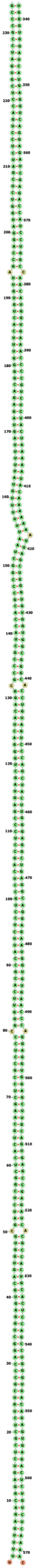
Long & highly paired dsRNA in the UTR of ZNF426. Structure was predicted using dsRNAscan and visualized here in <u>FORNA</u>. The structure was 312 bp long with only 9 mismatches and a single bulge, with dsRNA arms separated by a 194 nt loop (not shown).

**Figure S3.**
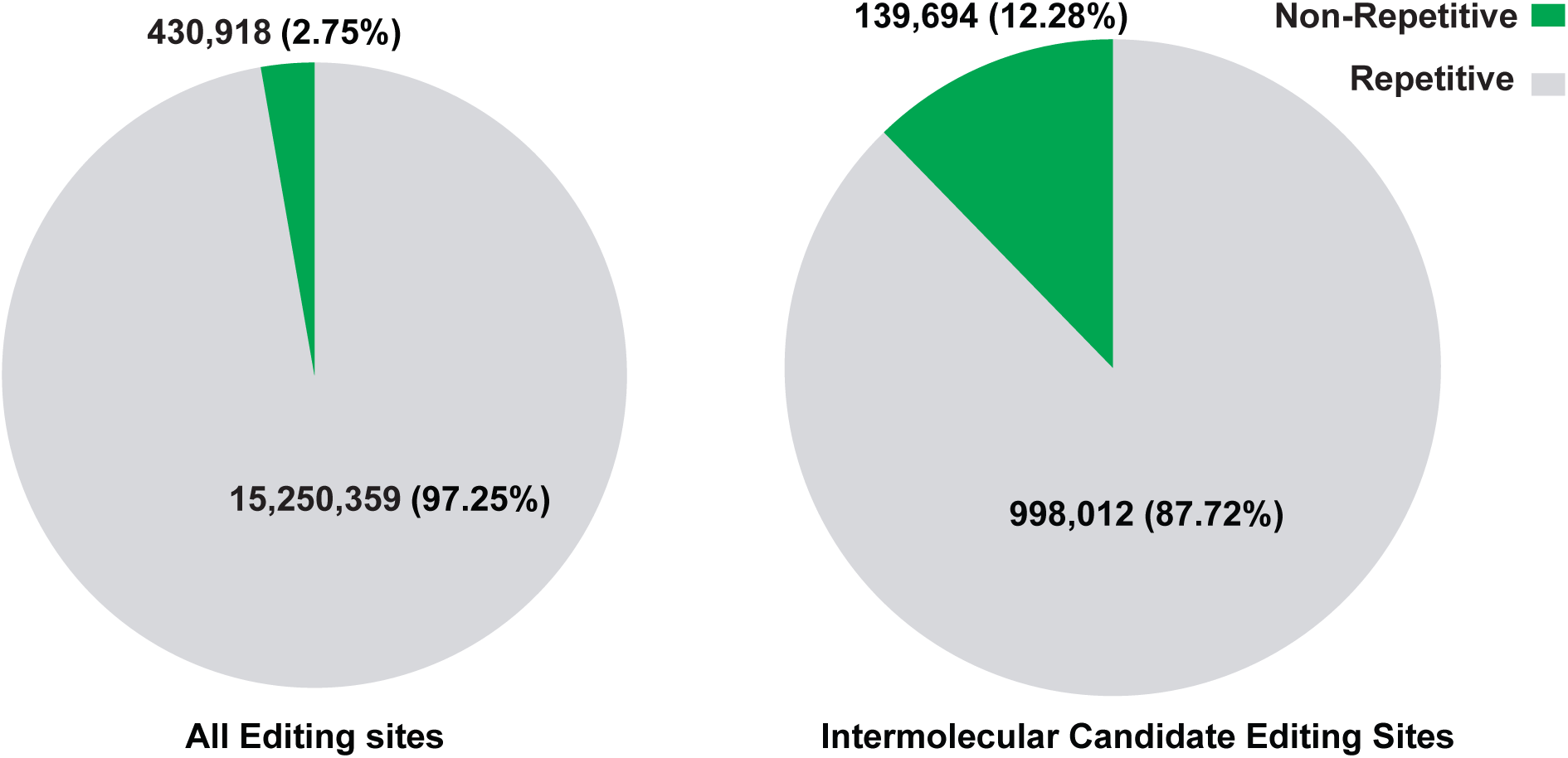
Comparison of non-repetitive editing sites. The total number of RNA editing sites (∼15.7 million) was obtained from the REDIportal database. These sites were then annotated based on whether they overlapped with repetitive elements from the Dfam database. The pie chart on the left shows that only 2.75% (430,918 sites) of all editing sites were found in non-repetitive regions, while the vast majority (97.25%, 15,250,359 sites) overlapped repetitive elements. On the right, the intermolecular candidate editing sites—defined as those that did not overlap any of our 5 million predicted dsRNA regions—were analyzed. Of these, 12.28% (139,694 sites) were non-repetitive, while 87.72% (998,012 sites) overlapped repetitive elements.

**Figure S4.**
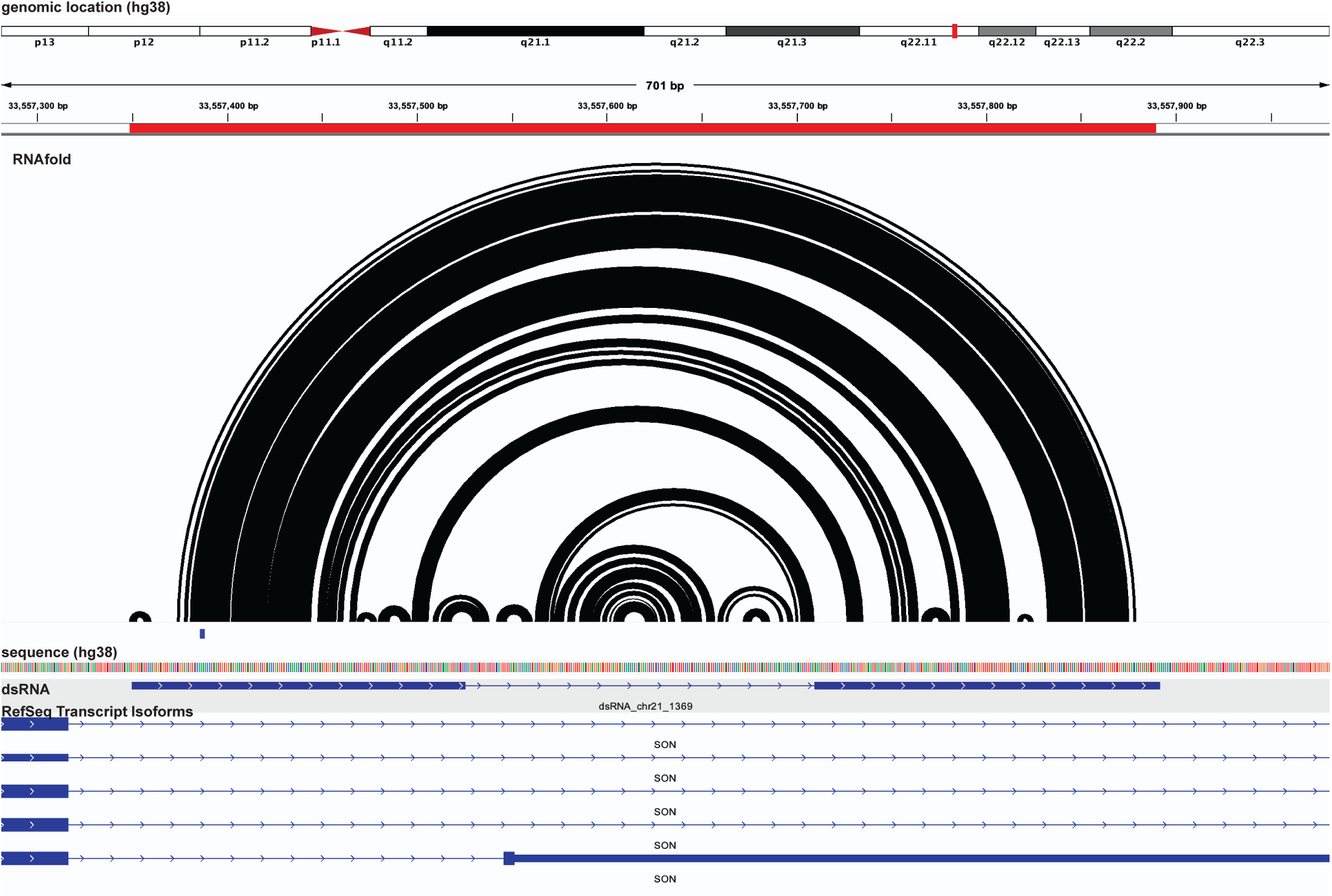
SON dsRNA location in IGV. This figure shows the genomic location and predicted base-pairing structure of the SON dsRNA, as predicted by RNAfold. The concentric arcs represent the RNA secondary structure, illustrating base pair interactions within the sequence. Below, the IGV browser view shows the alignment of the predicted dsRNA (dsRNA_chr21_1350) with the RefSeq transcript isoforms of the SON gene. The dsRNA is located in a completely intronic region for most SON isoforms shown, but in one isoform (bottom isoform), the dsRNA spans both an intron and the 3′UTR.

**Figure S5.**
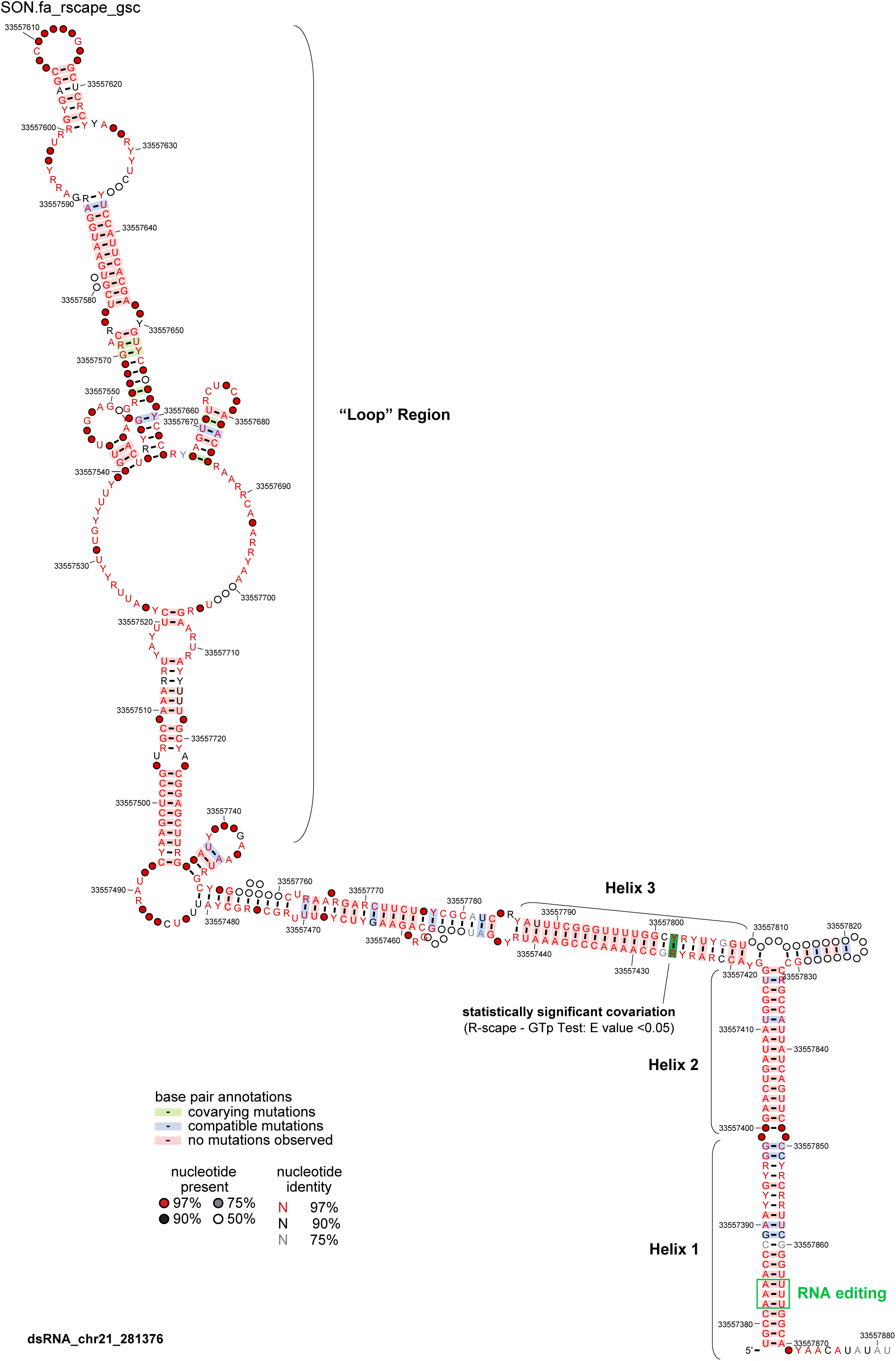
Consensus structure of the conserved dsRNA in SON. The structure consists of three helices and a loop region. The first three helices were initially predicted by dsRNAscan, while the loop region was added during alignment and conservation analysis (see Methods). Base pairs are color-coded and shaded according to covariation and mutation types, as shown in the legend. Covarying mutations from the alignment are shaded in red and green (via R2R) but do not reach statistical significance. The statistically significant covarying pair detected by R-scape (E-value < 0.05) is highlighted with a dark green box. Nucleotide identity across species is indicated by color and letters, following the legend. The three RNA editing sites within Helix 1 are marked with green outline. For detailed sequence alignment and structure prediction methods, see Methods.

**Figure S6.**
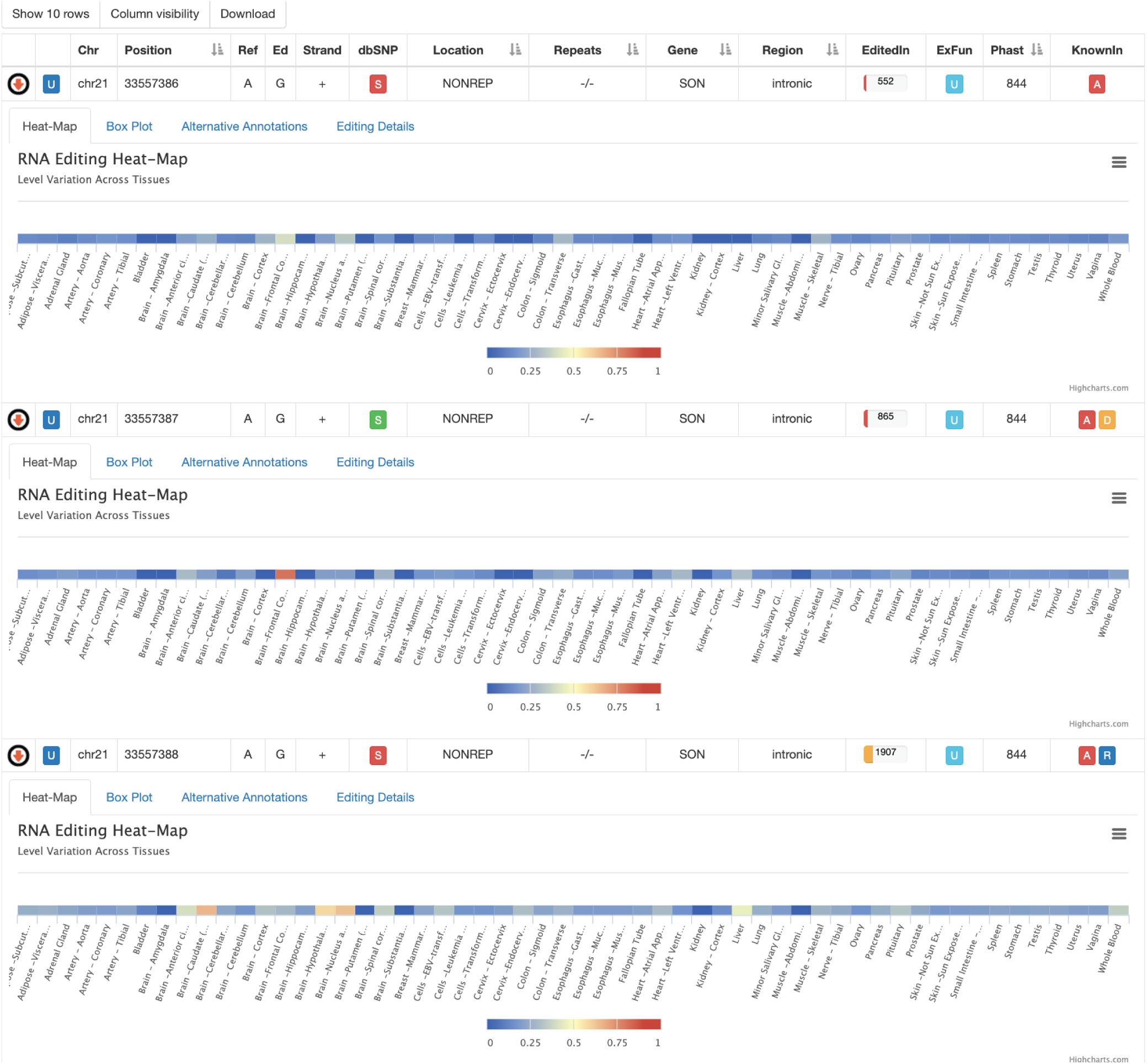
Editing sites in the SON conserved dsRNA. RNA editing levels at three specific A-to-G editing sites located within the conserved dsRNA structure in the SON gene, as retrieved from the REDIPortal database. All three sites are found in the intronic region and are non-repetitive (NONREP), with editing levels varying across different tissues. The heatmaps illustrate RNA editing frequency across 50 GTEx tissues, with blue representing lower editing frequency and red indicating higher editing frequency, where values range from 0 (no editing in any sample) to 1 (editing in all samples). These heatmaps highlight which tissues have the highest RNA editing levels, with the third site (chr21: 33557388) showing the most extensive editing across tissues. This site exhibits the highest editing index (1907), followed by the second (865) and first (552) sites. The variation in editing frequency emphasizes tissue-specific patterns of RNA editing in these conserved dsRNA regions.

